# Relevance learning via inhibitory plasticity and its implications for schizophrenia

**DOI:** 10.1101/161331

**Authors:** Nathan Insel, Blake A. Richards

## Abstract

Symptoms of schizophrenia may arise from a failure of cortical circuits to filter-out irrelevant inputs. Schizophrenia has also been linked to disruptions to cortical inhibitory interneurons, consistent with the possibility that in the normally functioning brain, these cells are in some part responsible for determining which inputs are relevant and which irrelevant. Here, we develop an abstract but biologically plausible neural network model that demonstrates how the cortex may learn to ignore irrelevant inputs through plasticity processes affecting inhibition. The model is based on the proposal that the amount of excitatory output from a cortical circuit encodes expected magnitude of reward or punishment (”relevance”), which can be trained using a temporal difference learning mechanism acting on feed-forward inputs to inhibitory interneurons. The model exhibits learned irrelevance and blocking, which become impaired following disruptions to inhibitory units. When excitatory units are connected to a competitive-learning output layer, the relevance code is capable of modulating learning and activity. Accordingly, the combined network is capable of recapitulating published experimental data linking inhibition in frontal cortex with fear learning and expression. Finally, the model demonstrates how relevance learning can take place in parallel with other types of learning, through plasticity rules involving inhibitory and excitatory components respectively. Altogether, this work offers a theory of how the cortex learns to selectively inhibit inputs, providing insight into how relevance-assignment problems may emerge in schizophrenia.

## Introduction

Many symptoms of schizophrenia can be understood as an inability of the brain to appropriately assign relevance to environmental stimuli and internal representations. Schizophrenic patients exhibit difficulties filtering-out, or gating, irrelevant external stimuli (Mcghie and Chapman, 1961; Venables, 1964; Lang and Buss, 1965; McGhie, 1970; Garmezy, 1977; Shagass, 1976; Venables, 1977; Garmezy, 1977), and delusions may also be the product of misattributing relevance (what others have labeled “salience”) to certain types of internally-generated representations (Kapur, 2003). Convergent evidence suggests this may result from a dysfunction of inhibitory processes. This idea may have been first proposed by Johnson (1985), who hypothesized that that schizophrenia symptoms arise specifically from a failure of feed-forward inhibition–i.e., activation of inhibition by a system’s inputs. Circuits for cortical feedforward inhibition are now relatively well defined, and may principally involve fast-spiking, parvalbumin-expressing (PV+) inhibitory interneurons (Agmon and Connors, 1992; Gibson et al., 1999; Dantzker and Callaway, 2000; Gonchar and Burkhalter, 2003). It is also now well established that PV+ interneurons are compromised in schizophrenia (reviewed by (Benes and Berretta, 2001; Lewis et al., 2005; Lewis and Moghaddam, 2006; Lewis, 2014; Gonzalez-Burgos et al., 2015)).

Computational models have helped to articulate the link between inhibitory dysfunction and schizophrenia (Vogels and Abbott, 2007, 2009; Murray et al., 2014). An important example is work by Vogels and Abbott (2007, 2009) which demonstrated how inhibition may serve to selectively gate some representations but not others. A theme of these models is the importance of balanced excitation and inhibition (EI balance) within the network. EI balance has been extensively studied across a range of cortical regions (e.g., auditory cortex (Wehr and Zador, 2003; Zhang et al., 2003; Dorrn et al., 2010), somatosensory cortex (Gabernet et al., 2005; Wilent and Contreras, 2005; Daw et al., 2007; Chittajallu and Isaac, 2010), olfactory cortex (Poo and Isaacson, 2009), visual cortex (Anderson et al., 2000; Xue et al., 2014), and frontal cortex (Haider et al., 2006)). EI balance may fluctuate dynamically, such as through disinhibition, the importance of which has gained increasing recognition (Pi et al., 2013; Isaacson and Scanziani, 2011; Carcea and Froemke, 2013; Letzkus et al., 2015; Kato et al., 2015). Therefore, a better understanding of the relationship between cortical disinhibition and relevance coding may be critical to understand schizophrenia.

In the present study, we sought to fill gaps in our understanding of the link between relevance assignment and neural inhibition. Two main questions are addressed. First, what may be the relationship between relevance-coding and the dynamic fluctuations in EI balance? Second, how might inhibitory components of the cortical circuit learn the relevance of specific input patterns? Answering these questions will help explain how “gating” is learned, and thereby also help to explain symptoms of schizophrenia. To this end, we have developed an abstract, but biologically plausible, neural network model that is capable of learning to ignore specific inputs, but not if inhibition is disrupted. The fundamental proposal in the model is that the overall level of excitation in a cortical circuit can be interpreted as a signal of the temporally discounted expectation of rewards and/or punishments (**Figure 1A**; see also Insel and Barnes 2015). According to this formulation, deviations in EI balance come to represent the network’s estimate of the magnitude of the value signal used in reinforcement learning (Sutton and Barto, 1998).

**Figure 1.**
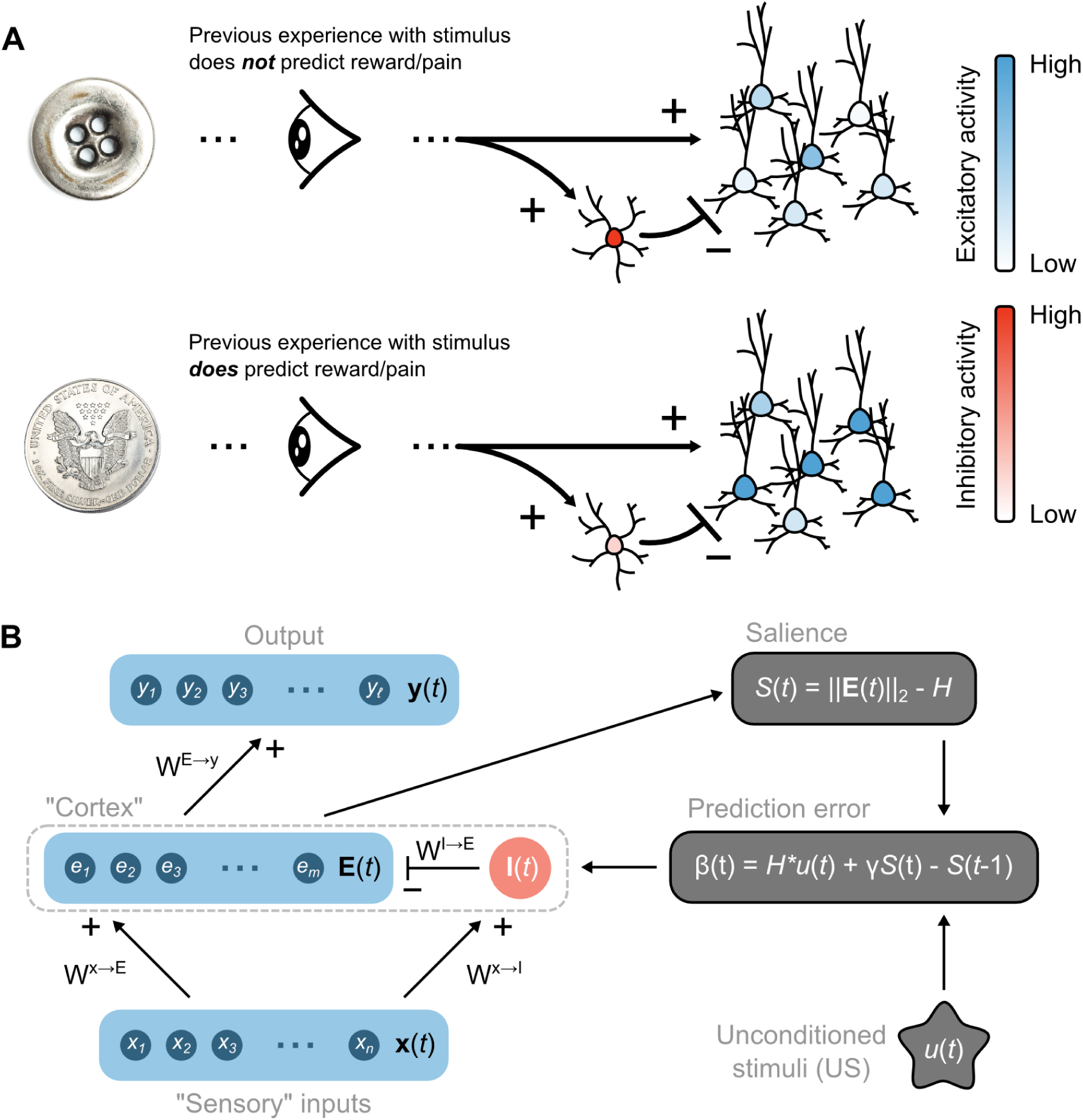
Overview of the proposed relevance code and network model. (**A**) Schematic illustrating the hypothesis that relevance (prediction of reinforcement) is coded by levels of excitatory neuron output from a network, which is controlled by feed-forward inhibition. (**B**) Basic structure of the network model. Left side shows feed-forward connections from “Sensory” inputs, through inhibitory (I(t)) and excitatory (E(t)) “Cortex” units, with E(t) units feeding onto an output layer. Right-side shows how the salience signal (S(t)), computed from the overall level of excitatory unit activity, is combined with signals about environmental unconditioned stimuli (u(t)) to generate a prediction error that supervises the plasticity of connection weights between Sensory and Cortex layers.

Three sets of simulations are used to demonstrate the explanatory power of the model. The first demonstrate the model’s capacity to learn input relevance/irrelevance, and that, paralleling symptoms of schizophrenia (e.g., (Jones et al., 1992, 1997; Oades et al., 2000)), learning is disrupted by impaired inhibition. The second set of simulations use an extended model to show how the proposed relevance code can modulate learning and, furthermore, reproduce the effects of manipulating inhibition in rodent frontal cortex (Piantadosi and Floresco, 2014; Courtin et al., 2014). The final set of simulations show how relevance learning could occur concurrently with other types of learning, e.g. learning to categorize input patterns, thereby providing a mechanism to multiplex information about stimulus-relevance and stimulus-identity.

## Methods

### Network summary

The basic structure of the neural network model is illustrated in **Figure 1B**. The core of the model is a spiking, two-layer feedforward neural network composed of different types of units. Stimuli are encoded by a set of input units, the ‘Sensory’ layer, **x**(*t*) = [*x*_1_(*t*), …, *x*_*n*_(*t*)] (for notation purposes, we use bold symbols for any variable that represents a population of neurons in the model). This layer is a set of excitatory Poisson units with firing rates ***ψ***(*t*) = [*ψ*_1_(*t*),*…,ψ*_*n*_(*t*)]. Changes in the Sensory layer take place when stimuli are presented, as described in more detail below. Sensory units feed into a middle layer, ‘Cortex’ that is comprised of two populations: an excitatory population of linear-non-linear-Poisson units (Paninski et al., 2007) with rates ***λ***(*t*) = [*λ*_1_(*t*),…,*λ*_*m*_(*t*)], and spiking activity **E**(*t*) = [*e*_1_(*t*),*…,e*_*m*_(*t*)], and a single inhibitory population unit, **I**(*t*), which receives **x**(*t*) and inhibits the excitatory neurons divisively. The receptive fields of Cortex units have no temporal dimension, so the activity at any point only reflects the current inputs to the network. The inhibitory unit is intended to model the entire population of inhibitory interneurons that provide feed-forward inhibition in the circuit. We justify this simplification in part by noting evidence that feed-forward inhibition in cortex provides a non-specific “blanket” of inhibition to excitatory neurons (Fino et al., 2013; Karnani et al., 2014).

The model is highly abstract, and makes a number of simplifying assumptions for the sake of mathematical tractability. One notable simplification, in comparison to true cortical networks, is the lack of feedback connections between excitatory units. The decision to omit these eliminated the need to explicitly model feedback inhibition to balance recurrent excitation, which helped focus the present investigation on the hypothesis that plasticity in feed-forward inhibition can support relevance learning. A second notable simplification is the absence of explicit signaling loops through other structures (e.g. the basal ganglia, thalamus, or other layers of cortex), which may be necessary for relevance learning in the brain. Naturally, the computations illustrated by the present, simplified network may be implemented in the brain through more steps and across more regions. Ultimately, the model serves as a proof of concept that speaks to only certain elements of true cortical computation.

The connections from input units to the excitatory cortex units are contained in an *m × n* synaptic weight matrix, *W*^**x**→**E**^, the connections from input units to the inhibitory Cortex unit are contained in the *n*-dimensional vector of synaptic weights, *W*^**x**→**I**^, and the connections from the inhibitory unit to the excitatory units are contained in an *m*-dimensional vector of synaptic weights, *W*^**I**→**E**^.

Altogether, this set-up gives the following equations which describe the activity of the model:

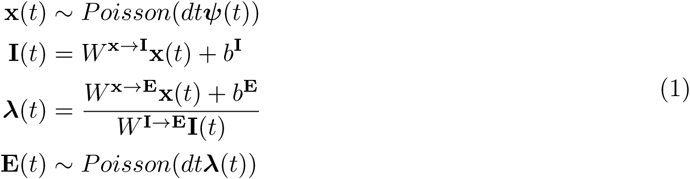

where *dt* is the time-step, which is 20 ms for most simulations, and *b***^E^** and *b***^I^** are bias terms.

One additional component that is not included in the above equations, but which contributes to relevance learning (see Relevance Learning, below), is a signal communicating the unsigned magnitude of reward or punishment, i.e. the unconditioned stimulus (US). In the present simulations the value of the US at a given time (u(t)) is either 1 or 0, though in principle it could as easily be a graded value with 1 representing the highest degree of relative salience and 0 the lowest.

In some simulations, we add an additional output layer of units with activity **y**(*t*) = [*y*_1_(*t*), *…,y*_*ℓ*_(*t*)] that receives inputs from the excitatory cortical units via an *ℓ × m* synaptic weight matrix, *W* ^**E**→**y**^. In those simulations which address the effects of frontal manipulations on fear behavior, the output layer is intended to represent an amygdala (’Amygdala’) and implements a competitive learning algorithm (according to the framework of Rumelhart and Zipser 1985). In the competitive learning module, a maximum of only one unit may be active at any given time (it is possible for no units to be active). Whether a unit, *i*, is active depends on two conditions: (1) the unit is receiving stronger input than any of the other units, (2) the unit’s input, *z*_*i*_(*t*) is greater than a threshold *θ*. Amygdala units also receive signals from the US, such that *u*(*t*) can help to increase input, *z*_*i*_(*t*). Based on all of this, the activities of the Amygdala units are governed by the following equations:

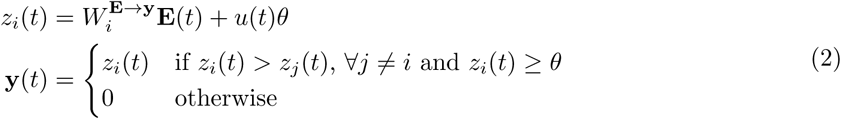

where 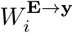 is the vector of synaptic weights into the *i*_*th*_ Amygdala unit. There are some important aspects of this to note:

1. All units will pass threshold, *θ*, if *u*(*t*) = 1.
2. Every 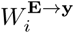 vector has its magnitude normalized such that 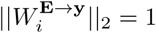 (similar to Rumelhart and Zipser 1985).
3. Only the unit whose weight vector 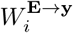 is most closely aligned to **E**(*t*) (according to the dot product) will be active.

The threshold, *θ*, determines when the Amygdala layer can have any active units in the absence of an unconditioned stimulus (i.e. if *u*(*t*) = 0). An explanation for how *θ* was derived mathematically is given below in the Amygdala learning section.

Although having a single neuron firing is undoubtedly not what occurs in the mammalian amygdala, there is evidence for a competitive “winner-takes-all” mechanism (Han et al., 2007; Rashid et al., 2016), such that a single ensemble of neurons is active and all others are silent. Therefore, the active unit in our model ‘Amygdala’ could be taken to represent an ensemble of “winning” neurons. Since individual units in this case were representative of larger ensembles, the winning unit’s firing rate was kept as a continuous ”activation level” value rather than Poisson-distributed spiking.

The output layer takes on a different form in those simulations that demonstrate how our model can multiplex relevance signals and stimulus identity. In this case, the output units represent some efferent, such as a second area of cortex, that is responsible for categorizing input activity. For simplicity, we refer to this layer in the simultations as the ‘Category’ layer. The Category layer is a set of softmax, linear-non-linear-Poisson units with rates *ϕ* (*t*) governed by:

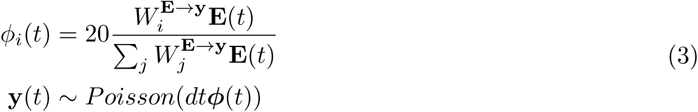

where the activity of the units is proportional to the probability of each of the *𝓁* possible categories for the current stimulus (a rate-of-fire of 20 Hz corresponds to a probability of 1).

### Stimuli

Sensory (input) units are divided into sets of stimulus-coding and non-coding units. Each stimulus is capable of activating one tenth of the Sensory units. In the case of the Learned Irrelevance simulation, for example (see Learned irrelevance and blocking, below), one tenth of the units are activated by the stimulus, ”CS+” that is paired in time with the US, one tenth of the units are activated by another stimulus, ”CS-”, that is not paired with the US, and the rest are activated by neither CS+ nor CS-. Importantly, Sensory units fire both when activated by a stimulus and not, just at different rates. An active sensory unit fires spikes with a Poisson process at a rate of *ψ*^*on*^ = 20 Hz, while an inactive sensory unit fires spikes with a Poisson process at a rate selected randomly based on a gamma distribution that peaks at 0.6 Hz and has a variance of 3 Hz. These rates were selected based on baseline firing characteristics among putative excitatory neurons recorded from the rat medial prefrontal cortex (Insel and Barnes, 2015). Hence, for example, if the CS+ is presented to the network then the ten percent of the units activated by CS+ will be firing at a rate of 20 Hz, while other units will continue firing at their typically low (0-2 Hz) but occasionally high (10 or 20 Hz) baseline rate. In those simulations that use the Category output layer, Sensory units are divided into ten sets, with each set activated differentially by a particular category (in these simulations, baseline rates were also simplified to be homogeneously 2 Hz, as variance was found to not impact the results).

### Weights

Initialization of the three sets of connection weights–Sensory to Cortex excitatory units (*W* ^**x**→**E**^), Sensory to Cortex inhibitory unit (*W* ^**x**→**I**^), and Cortex inhibitory to excitatory units (*W* ^**I**→**E**^)–took into account three issues. First, when novel stimuli were first presented to the network, the evoked activity in Cortex excitatory units needed to be higher than baseline levels, but not higher than levels associated with “relevant” stimuli (described below in Relevance learning). Second, baseline activity of the cortex inhibitory unit had to be high enough that reducing *W* ^**x**→**I**^ had an impact on Cortex excitatory units. Third, that *W* ^**x**→**I**^ were balanced with *W* ^**x**→**E**^, such that small changes in *W* ^**x**→**I**^ could not dramatically alter population activity.

Starting weights were assigned as follows: all inhibitory to excitatory weights (*W* ^**I**→**E**^) were set to 0.4. Sensory to inhibitory weights (*W* ^**x**→**I**^) were set to 0.5, although in most simulations these were adjusted during pre-exposure periods to establish a desired level of Cortex activity. In the initial simulations which did not make use of an output layer, Sensory to Cortex excitatory unit weights (*W* ^**x**→**E**^) were selected randomly from a Gaussian distribution centered at 0.1, with a standard deviation of 0.4, and any negative connections were clipped at zero. This yielded an average weight of 0.2, with many at zero. In those simulations which used the Amygdala output layer, fear learning to the CS+ required that Sensory activity patterns could be interpreted from Cortex excitatory unit patterns; therefore, Sensory to Cortex excitatory weights (*W* ^**x**→**E**^) were set to a Gaussian-smoothed identity matrix (the smoothing window was equal to the number of inputs divided by 20; e.g., for 500 units, a window of 25 was used), then multiplied by 1/3 the number of inputs. In those simulations which used the Category output layer, Sensory to Cortex excitatory weights were modified according to stimulus presentations; initialization weights were therefore arbitrarily drawn from a uniform distribution over [0 − 1].

### Relevance Learning

The principal learning mechanism used in this paper is a modification of the temporal difference learning algorithm (Sutton and Barto, 1998). Specifically, a population-based relevance (or “salience” signal), *S*(*t*) reflects the deviations in excitatory activity from an established baseline. The baseline level can be thought of as the EI balance set-point maintained by the cortical network. The level of Cortex excitatory unit activity was measured as the vector norm of the population of spiking excitatory units, 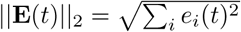(the reasons for using the norm become clear in the following section on Amygdala learning). *S*(*t*) is therefore determined by the difference between ‖E(*t*)_2_‖ and the homeostatic set-point for the population, *H* (**Figure 1B**):

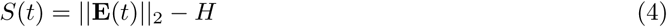

The goal of relevance learning in the network is to have *S*(*t*) come to represent expected relevance, which we interpret as “unsigned value”, *U* (*t*):

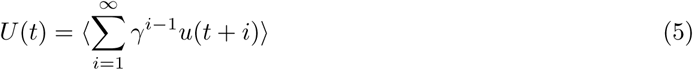

where *u*(*t*) is the unsigned reward/punishment signal, US, described above, *γ* is a temporal discounting factor, and ⟨.⟩ indicates expected value. *U* (*t*) is akin to the value function, *V* (*t*), used in temporal difference learning (Sutton and Barto, 1998). Similar to temporal difference learning, the goal of learning in our model is, in part, to ensure that *S*(*t*) is a good estimate of *U* (*t*). This is accomplished using a prediction error signal, *β* (*t*):

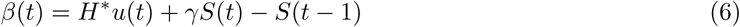

where *H*^*\**^ is a salience scaling factor that determines how much cortical activity levels should deviate from the set point in response to relevant stimuli. We use *β* to represent our prediction error signal, rather than the usual *δ*, to distinguish it from prediction error signals that measure differences in signed (as opposed to unsigned) value estimates (Sutton and Barto 1998); the notation also deviates slightly from convention by using ”t” rather than ”t+1”, to avoid questions about whether the model has future information).

To put it another way, the system learns to ensure that fluctuations in ∥**E**(*t*)∥_2_ away from the set-point, *H*, reflect experience with rewards/punishments (the US). The scale of the fluctuations is determined by *H*^*\**^. Training the salience signal *S*(*t*) involves updating the synaptic weights in Cortex to achieve *β*(*t*) = 0. It can be seen that this is achieved when:

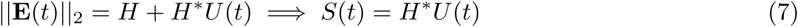

since *U* (*t* − 1) = ⟨*u*(*t*)⟩ + *γU*(*t*). Therefore, *β* is generally close to zero when the following conditions have been achieved: (1) for stimuli that do not predict any reward or punishment, the norm of the spike count in the excitatory cortical population is equal to the homeostatic constant, *H* and (2) for stimuli that do predict reward or punishment *S*(*t*) is a linear function of *U*(*t*) with a slope of *H*^*\**^. Once again, the highest possible US (*u(t)*) value is 1 based on the assumption that natural stimuli, no matter how salient, should not be capable of pushing network activity beyond a limit.

To learn this, we apply a Hebbian weight update modulated by the prediction error, *β* (*t*). In most simulations the weight change takes place on the feed-forward connections to the inhibitory unit, described by the following update equation:

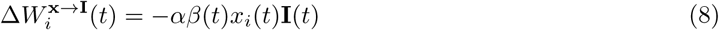

where Δ indicates a weight update, *i* indexes sensory units, α is the learning rate, and the negative sign is required because the weight change is on synapses that have an overall inhibitory effect. The synapses were updated on-line at each time-step of the simulations.

In early simulations, the performance of this learning rule is compared against modified rules in which either the *W* ^**x**→**E**^ or *W* ^**I**→**E**^ synapses are modified:

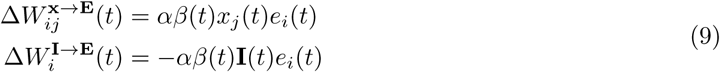

where *j* refers to cortex excitatory units.

An important question raised by the above relevance-learning equations is how real cortical networks implement a homeostatic set-point, and what may account for its deviation during rewards/punishers. We offer two non-exclusive possible conceptualizations. The first follows from a tradition of research on prediction error signals in neuromodulatory systems like dopamine and norepinephrine, the second builds off of recent research on the mechanisms by which EI balance is maintained within cortical networks. In the former scenario, the salience signal is explicitly read out by cells in neuromodulatory nuclei, as has been described in dopaminergic nuclei (Bromberg-Martin et al., 2010), which is then used to compute a prediction-error signal that feeds back to the cortical afferent, modifying plasticity (the Hebbian weight update) accordingly. In the second scenario, the prediction error calculation responsible for maintaining EI balance are local to the cortex, and may take place, for example, by intrinsic signaling processes within inhibitory neurons in response to feedback from local excitatory neurons (i.e. the salience signal, see Vogels et al. 2011; Luz and Shamir 2012). Either the salience signal itself (in this scenario, the inhibitory neuron responses to excitatory signals) or the cellular processes responsible for computing the difference between those signals and the desired level (the prediction error signals) are modified by neuromodulatory signals carrying information about current rewards/punishers. The present model is built at one level of abstraction above this implementation, remaining agnostic as to where and how the signals are implemented. These details are, however, likely to be valuable for future empirical and theoretical investigations (this is also considered in Discussion).

### Amygdala learning

In simulations with output units, **y**(*t*), such as the Amygdala, synapses between Cortex excitatory units and the output units were also trained. In simulations using an Amygdala output layer (which were used to simulate the effects of frontal cortex manipulations on fear responses), the Cortex-to-Amygdala weights, 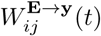, were trained with a competitive learning algorithm based on Rumelhart and Zipser (1985). More precisely:

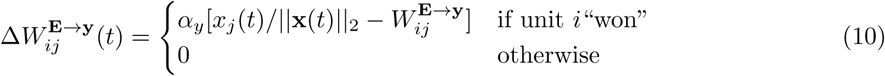

where *α*_*y*_ is the learning rate. These weight updates took place simultaneously with weight updates within Cortex for relevance learning.

Importantly, if a given unit in the Amygdala, *i*, always tends to “win” when presented with excitatory population vectors 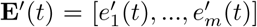 sampled from the rate vector, **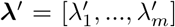**, then this learning algorithm will converge (on average), when:

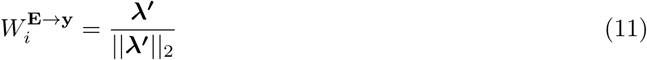

This implies that the weight vector 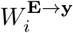 is actually just ***λ***ʹ scaled to have a norm of 1. As such, if the excitatory Cortex units generate activity vectors with consistent norms that scale based on previous associations with reward or punishment (which should be the case if relevance learning has converged, see equation (8)), then when Amygdala learning has converged the input to unit *i, z*_*i*_(*t*) (as determined by 2), will be the maximum in response to ***λ***ʹ. (That is because the dot product of two vectors is a function of the angle between them, with angles of zero giving the greatest dot product when the magnitude is constant). As such, a suitable threshold, *θ*, can be found to ensure that in the *absence* of a US the Amygdala only responds to stimuli that have been paired with reward or punishment in the past so that the Amygdala learning algorithm could converge. In the simulations presented here, the value of *θ* was set by grid search so that the probability of any neuron crossing threshold would be very low if no learning had occurred, and very high if a US was present or learning had converged.

### Categorization learning

In simulations where we trained the output units to categorize input stimuli, we used back-propagation of error (Rumelhart et al., 1986) to train the weight matrices *W* ^**x**→**E**^ and *W* ^**E**→**y**^. More precisely, target vectors, **o**(**x**(*t*)) are defined, where each stimulus provided to Sensory units has a corresponding target vector for the output Category units. The cross-entropy (Rumelhart et al., 1986) between the Category activity and target vectors was used as the loss function to train the network:

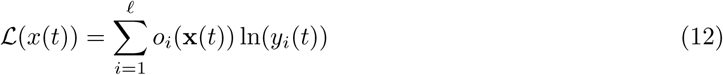

where *o*_*i*_(**x**(*t*)) is the “target” response to input **x**(*t*) for output unit *i*, i.e. *o*_*i*_(**x**(*t*)) = 1 if *i* is the correct category for **x**(*t*), and it is zero otherwise.

For any weight *W*_*ij*_ in *W* ^**x**→**E**^ or *W* ^**E**→**y**^, the weight update is determined by the partial derivative of this loss function with respect to the weight:

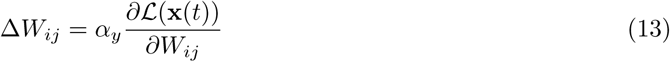

where *α*_*y*_ is the learning rate. This ensures that the network learns to correctly categorize the stimuli (i.e., the pattern of Sensory unit activity, **x**(*t*)) using the output, Category units **y**(*t*). As with Amygdala learning, the categorization learning proceeded in tandem with the salience learning.

### Learned irrelevance and blocking

The first set of simulations tested learned irrelevance and, in a separate protocol, blocking. These were both run using *dt* = 20 ms. At the beginning of the simulations, a 60 s adaptation period without stimulus presentations was simulated, which allowed weights to adjust to the randomly-selected baseline input activity levels. All pharmacological simulations were implemented after adaptation.

#### Learned irrelevance

Each of two stimuli used an independent, inter-trial-interval jittered between 20 and 30 seconds. Experimental stimuli were simulated as 200 ms periods during which the firing rate of a pre-determined set of Sensory units (10%) was raised to *ψ*^*on*^. One of the two stimuli, CS+, was paired with a reinforcer, or unconditioned stimulus (US) by setting *u*(*t*) = 1 (see the section Stimuli above). The onset of the CS+ preceded the US by 100 ms. The network was capable of learning the relevance of a temporally offset CS+ because *β*(*t*) integrated signals across timesteps with a discounting factor.

#### Blocking

The blocking protocol used the same parameters as that of learned irrelevance, with the exception that there were four phases of stimulus exposures: 1) a pre-exposure phase, in which both CSs (named CS-A and CS-B) were presented 50 times *without* the US (inter-trial-intervals were decreased to 10-15 s to reduce runtime), 2) a conditioning phase, in which CS-A was presented 50 times paired with a US, 3) a blocking phase, in which the CS-B was presented 50 times simultaneously with the CS-A, paired with a US, 4) a test phase, in which the CS-A and CS-B were presented independently 10 times in the absence of a US. To reduce runtimes, the inter-stimulus intervals were also reduced to between 10 and 15 seconds.

#### Tests of inhibitory disruptions in different model versions

The effects of inhibitory connection strength changes on learned irrelevance and blocking were assessed using different model versions. The model versions differed with respect to which synapses were plastic: *W* ^**x**→**I**^, *W* ^**x**→**E**^, or *W* ^**I**→**E**^, equations (8) and (9) above. The purpose of the test was to evaluate whether disruptions to inhibitory systems correspondingly disrupt learned irrelevance and blocking, as has been hypothesized to take place in schizophrenia. Inhibitory disruptions were made by reducing the degree to which excitatory units could respond to the inhibitory units by 10%:

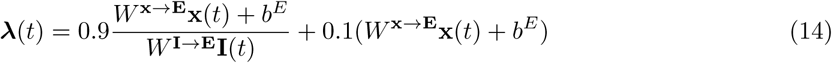

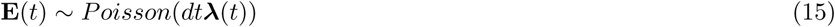

### Simulation of Experimental Data: Pharmacological Effects on Latent inhibition

Piantadosi and Floresco (2014) demonstrate the effect of GABA-A antagonists on latent inhibition. Latent inhibition refers to the classic behavioral phenomenon whereby it is harder to associate a familiar stimulus (one that a subject has been pre-exposed to) with a reinforcer (Lubow and Moore, 1959). Latent inhibition is also known to be disrupted in schizophrenia (Lubow et al., 7 06; Baruch et al., 1988; Rascle et al., 2001; Gray and Snowden, 2005). As shown in **Figure 4A-D**, a protocol was created that matched the one used in rats (see also McAllister 1997; Enomoto et al. 2011). For processing time purposes, the stimulus and inter-stimulus times used in the original were reduced by a factor of 5. The protocol began with a 60 s adaptation period, followed by three phases: 1) a pre-exposure phase, in which the network was presented with the conditioning stimulus (CS) 30 times (10% of input units, 6 s long, inter-stimulus interval of 6 s), 2) a conditioning phase, in which the CS was presented simultaneously with foot shock (*u*(*t*) = 1), and 3) a test phase, in which the CS was presented by itself 4 times. The protocol was performed on three pairs of network models, with each pair including one network given pre-exposures and one that was not given pre-exposures. The three pairs simulated the treatment groups used in the original study: animals treated with saline were simulated without any modification to the network, treatment with GABA-A antagonist during conditioning were simulated using a 20% reduction in inhibition, according to equation (14) during the conditioning phase, and treatment with antagonist during testing were simulated with the same disruption during the testing phase. Conditioned fear responses were measured as the maximal response of amygdala units, averaged across all timesteps during CS presentation.

### Simulation of Experimental Data: Optogenetic Effects on Fear Expression

Recent work by Courtin et al. 2014 found that inhibition of parvalbumin, fast-spiking neurons in the mouse mPFC can evoke fear responses, while excitation of the same neurons can decrease fear responses. The protocol used in that study was presently simulated as precisely as possible (**Figure 4E-H**), using all of the same parameters as used in the latent inhibition design. To stimulate optical stimulations, 20% reduction in inhibition, according to equation (14). During the pre-conditioning phase, this reduction in the inputs was applied for 250 ms intervals separated by 860 ms (equivalent to 0.9 Hz stimulations, as in the original study). This was followed by a conditioning phase, in which a 6 s CS+ was paired with footshock (i.e., the firing rate of input units coding for the CS was set to *ψ* ^*on*^ and *u*(*t*) = 1). As in the previous protocol, all stimulus and inter-stimulus times were decreased from the original study by a factor of 5. One change from the original protocol is that the 1 s US presentation used in the original study was lengthened to the entire CS period. We justify this change based on an assumed difference between real brains and the model: whereas in the brain, activity and plasticity are likely regulated by change, such as the onset or offset of a stimulus, the model treats each time point equivalently. Thus, the period during which the CS is on but US is off will extinguish the associations learned during their concurrence. The CS-US pairings were presented 12 times with an inter-trial interval of 4-30 s. The conditioning phase was followed by an extinction phase, in which the CS was again presented 12 times with the same inter-trial interval, followed in turn by a series of CS presentations accompanying the 40% reduction in *W* ^**x**→**I**^. To test the effect of inhibitory activation during a conditioned CS, the same conditioning protocol was used, but was followed by presentations of the CS accompanying increases to inhibitory unit activit. We found that only a 10% increase in *W* ^**I**→**E**^ was necessary to elicit changes approximating those observed in the original study.

### Pairing of relevance-learning with classification learning

To examine the ability of the network to carry both the salience signal and the other information simultaneously (i.e. to multiplex the salience signal with other signals) simulations were run wherein the feed-forward excitatory weights (*W* ^**x**→**E**^ and *W* ^**E**→**y**^) were trained to perform categorization of the inputs, **x**(*t*), while the excitatory weights onto the inhibitory unit, *W* ^**x**→**I**^, were trained according to the relevance algorithm described in equation (8). To do this, stimuli were presented as described above for 1000 timesteps with *dt* = 20 ms, and the learning proceeded at each timestep as described in Categorization learning, above. For simplicity, however, the network was trained using a batch method, wherein all the weight updates in response to all of the stimuli were calculated and applied simultaneously.

## Results

### Inhibitory interneuron plasticity can mediate learned irrelevance

The first simulations established that the model could learn to distinguish relevant versus irrelevant stimuli based on whether or not the stimuli were reinforced. The network was presented with two independent, 200 ms long stimuli with inter-trial-intervals that were independently jittered between 20-30 seconds (**Figure 2A**, *colored boxes on bottom*). The conditioned stimulus (CS+) was paired with an unconditioned stimulus (US) that began 100 ms after CS onset and terminated 100 ms following offset; the second stimulus was not paired with a US (CS-). Initially, the increased input from the Sensory layer, **x**(*t*), associated with presentation of either stimulus evoked an increased response among Cortex excitatory units (**E**(*t*)) and, by extension, the population-based salience signal (*S*(*t*); **Figure 2A**, *left*). As the number of presentations accumulated, there was a selective reduction in the degree to which excitatory units responded to the CS-, but no reduction in the degree to which the network responded to the CS+ (**Figure 2A**, *right*). **Figure 2B** illustrates the gradual decrease in excitatory unit population responses to the CS-(*left*) and the corresponding increases in the Cortex inhibitory unit (**I**(*t*)) response (*right*).

**Figure 2.**
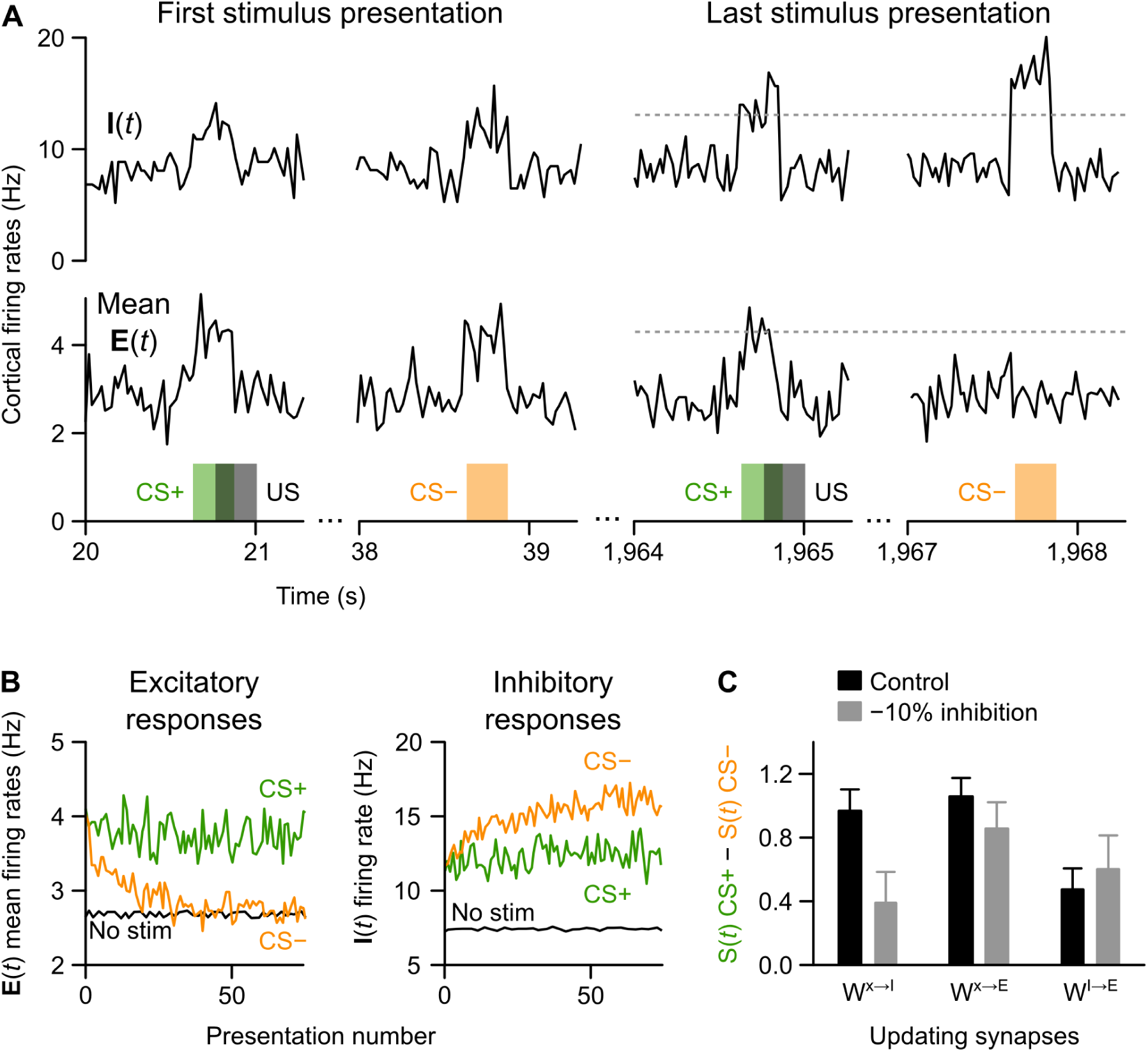
Demonstration of learned irrelevance and its impairment following disruption to inhibition. (**A**) Average Cortex excitatory unit activity (lower plots) and inhibitory unit activity (upper plots) at simulated, 20 ms time steps in response to unlearned stimuli (left side) compared with the end of a series of repeated presentations (right side). Stimuli are either paired (CS+, red bar) or unpaired (CS-, blue bar) with an unconditioned stimulus (US, grey bar). Excitatory responses were initially high to both stimuli, but after learning they increased only in response to the CS+, demonstrating learned irrelevance for the CS-. (**B**) Averaged excitatory unit (left) and inhibitory unit (right) responses to the CS+ (red) and CS-(blue) across presentations, as compared with non-stimulus periods (grey line). Learned irrelevance took place over the first 20 trials, after which excitatory responses to the CS-plateaued to the same level as non-CS inputs. This was due to increased inhibitory responses to the CS-. (**C**) Excitatory responses to the CS+ relative to the CS-during final presentations are plotted for both control conditions and in simulations of inhibitory dysfunction (means *±* STD across 30 model runs). Learned irrelevance was lost with inhibitory neuron disruption only in the inhibitory neuron plasticity model (*W* ^**x**→**I**^).

The increased responsiveness of the Cortex inhibitory unit to the CS-over presentations was due to the gradually increased connection weights between the units of the Sensory layer and the Cortex-inhibitory unit (*W* ^**x**→**I**^), caused by the temporal difference learning rule. We next examined whether the same patterns could be observed using other model versions, in which synapses either between Sensory and Cortex-excitatory units (*W* ^**x**→**E**^), or between Cortex-inhibitory and excitatory units (*W* ^**I**→**E**^) were modified. This comparison included an examination of how the model responded to disrupted inhibition (see Methods). The results of these tests are described in **Figure 2C**. Both *W* ^**x**→**I**^ plasticity and *W* ^**x**→**E**^ plasticity models exhibited much greater ability to learn to distinguish relevant versus irrelevant stimuli, indicated by the salience signal (*S*(*t*)) during the CS+ relative to CS-. Both of these models were better at distinguishing CS+ and CS-than the *W* ^**I**→**E**^ plasticity model. However, disrupted inhibition only eliminated learned irrelevance in the *W* ^**x**→**I**^ plasticity model. Differences in how the model types responded to disrupted inhibition could be assessed statistically: even ten repetitions of the simulation was *×* more than sufficient to demonstrate an interaction effect between model type and inhibitory manipulation (two-way ANOVA, type × manipulation: *F*_(2,54)_ = 21.85, *p* = 10^−^07; one-way ANOVA comparing the disrupted inhibition conditions: *F*_(2,27)_ = 21.65, *p* = 2*x*10^−^06, multiple comparisons between all groups significantly different using a Bonferroni correction).

If we take as an assumption that learned irrelevance impairments found in schizophrenia or other disorders arise from changes in inhibitory neurons, consistent with observed pharmacological effects in rodents, then the patterns observed in the *W* ^**x**→**I**^ plasticity model offer the best fit to the pathology. This supports the hypothesis that relevance learning is mediated by inhibitory interneuron plasticity.

### Inhibitory interneuron plasticity can explain blocking data

Another well-established relevance learning phenomenon is ’blocking’, in which one stimulus that has been previously reinforced can occlude learning for another reinforced stimulus Kamin, L J. Blocking is also known to be affected in schizophrenia (Jones et al., 1992, 1997; Oades et al., 2000). To examine whether the model exhibited blocking, a standard blocking protocol was simulated, illustrated in **Figure 3A**. Two different stimuli were presented to the network, CS-A (non-blocked) and CS-B (blocked). The difference between the non-blocked, CS-A, stimulus and the blocked, CS-B, stimulus is that CS-A was conditioned alone with the US (following habituation pre-exposures) while CS-B was conditioned only when paired with CS-A (following pre-exposures and CS-A conditioning). When this type of protocol is used in either rodent (e.g., Kamin, L J; Ganesan and Pearce) or human (reviewed by De Houwer et al.) experiments, it leads to CS-A being recognized as relevant for reward/punishment, but CS-B being judged irrelevant. The blocking effect was measured in the model by comparing the response of the excitatory unit population to CS-A and CS-B during the final test sessions (**Figure 3B**, also **Figure 3C** *inset*). As predicted, the model exhibited the basic blocking effect seen in people and animals, with CS-A generating a large excitatory response and CS-B generating a small one (**Figure 3C**, *left inset*). Because learning was supported by inhibitory neuron plasticity (in this, the *W* ^**x**→**I**^ plasticity model), changes in population responses to both CS-A and CS-B over stimulus presentations paralleled changes in inhibitory neuron, **I**(**t**) responses (**Figure 3C**, *right inset*). Notably, a strong increase in inhibitory neuron activity was observed during the “blocking” phase of conditioning, reflecting feed-forward inhibition compensating for both stimuli being presented simultaneously (**Figure 3C**, *presentations 0-50*). We note that this is also consistent with observations of increased fast-spiking, narrow-waveform neuron activity during stimulus presentations and movement (Insel and Barnes, 2015).

**Figure 3.**
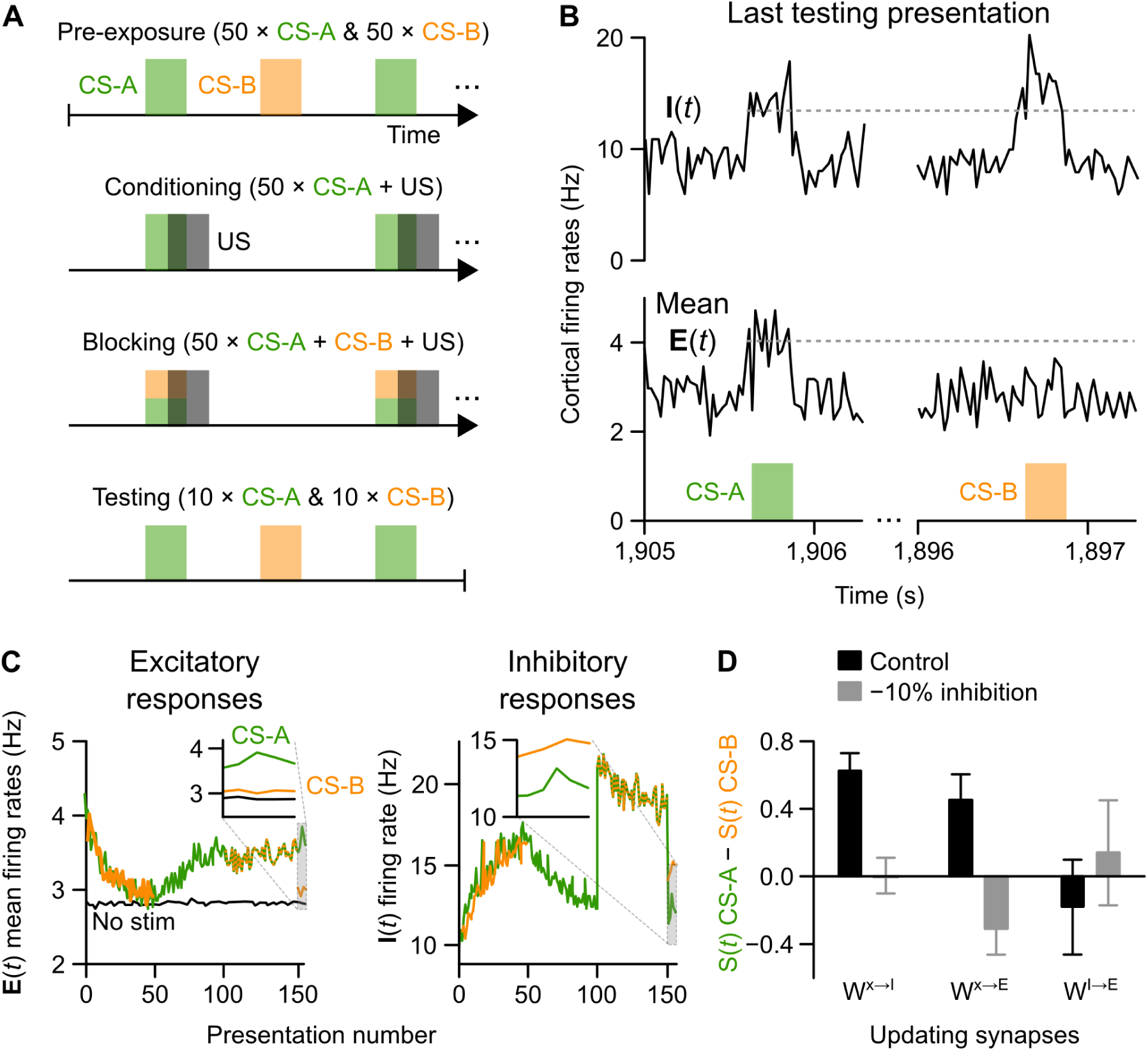
Demonstration of blocking and its impairment following inhibitory disruptions. (**A**) Illustration of the blocking paradigm: the model was first habituated to two stimuli (CS-A, CS-B; Pre-exposure), the CS-A and a US were then repeatedly presented at partially overlapping times (Conditioning), both CS-A and CS-B were then presented with the US (Blocking), followed by independent presentations of CS-A and CS-B (Testing). (**B**) Excitatory (lower plots) and inhibitory (upper plots) unit activity over 20 ms bins show the networks response to CS-A (left) and CS-B (right) at the end of the blocking paradigm. In spite of CS-B being paired with the US, the “blocked” stimulus did not elicit increased activity among excitatory units. (**C**) Excitatory (left) and inhibitory (right) unit responses to CS-A and CS-B over trials. Test epochs are expanded in insets. (**D**) Excitatory responses to CS-A relative to CS-B at the end of the test epoch are plotted in control simulations and simulations in simulations with dysfunctional inhibition (means*±* STD across 30 model runs). The inhibitory neuron plasticity model (*W* ^**x**→**I**^) showed a loss of the blocking effect when inhibition was disrupted; unexpectedly, the excitatory neuron plasticity model (*W* ^**x**→**E**^) exhibited a reversal of the blocking effect; i.e., CS-B was learned more strongly than CS-A.

As with the learned irrelevance simulation, blocking and the effect of inhibitory disruptions was tested in different versions of the models, defined by which synapses (*W* ^**x**→**I**^, *W* ^**x**→**E**^, *W*^**E**→**I**^) were plastic. Consistent with predictions, the blocking effect was eliminated in the *W* ^**x**→**I**^ plasticity model, following even a 10% disruption of inhibition (**Figure 3D**, *left bars*). Blocking was also observed in the *W* ^**x**→**E**^ model; however, in this model version an unexpected “reverse blocking” effect was observed following inhibitory disruptions (**Figure 3D**, *middle bars*). This was likely due to the supervised-learning mechanism becoming over-active following inhibitory disruptions, leading to a reduction in synaptic weights corresponding with CS-A presentations (regardless of it being paired with the US). No blocking effects could be obtained in the *W* ^**I**→**E**^ model (**Figure 3D**, *right bars*). Once again, these results could be judged statistically, with 10 repetitions more than sufficient to reveal an interaction between model type and manipulation (two-way ANOVA, *F*_(2,54)_ = 21.85, *p* = 10^−^07), one-way ANOVA of disrupted inhibition condition: *F*_(2,27)_ = 21.65, *p* = 2*x*10^−^06, with multiple comparisons test with Bonferroni correction showing a significant difference between *W* ^**x**→**I**^ and *W* ^**x**→**E**^ models).

Consistent with the results from the learned irrelevance simulations, the pattern of effects in the blocking paradigm, when considered in the context of schizophrenia symptoms and inhibitory neuron pathologies, lend credence to the hypothesis that blocking is mediated by plasticity in inhibitory neurons.

### Inhibitory relevance-learning network with amygdala module recapitulates effect of pharmacological manipulations on latent inhibition

Motivated in part by the pathology of schizophrenia, researchers have previously sought to experimentally link EI balance in frontal cortex to various forms of conditioning (Piantadosi and Floresco, 2014; Courtin et al., 2014). A set of simulations was designed to determine whether our model could recapitulate the findings of these studies. To explicitly simulate the link between experimental manipulations of the mPFC and behavior, it was necessary to demonstrate how the output of the mPFC, and in particular the relevance signal, *S*(*t*), might affect an efferent region that directly controls behavioral output. Since many of the rodent studies in this area use fear conditioning, the output layer was designed to represent the mamalian amygdala, and the levels of ’Amygdala’ unit activity were equated with fear expression. The Amygdala layer was modeled as a competitive network with 10 units receiving input directly from Cortex excitatory units (see Methods).

The first simulations examined the findings of Piantadosi and Floresco (2014), which showed that a GABA-A receptor antagonist, applied to the medial prefrontal cortex (mPFC), can have different effects on latent inhibition when applied at different phases of the learning protocol. As described above, latent inhibition refers to the phenomenon wherein it is harder to associate a stimulus with a reinforcer if a subject has previously been exposed to that stimulus. In the study by Piantadosi and Floresco (2014), animals were separated into two groups: those that received pre-exposures of the CS (PE) and those that did not (NPE). When the CS was subsequently paired with a footshock, the PE group was less likely to learn the fear association compared with the NPE group (i.e. the animals exhibited latent inhibition). Importantly, the authors found that blocking GABA-A receptors had different effects if done during the conditioning period or during the test: GABA-A antagonists infused during conditioning amplified latent inhibition, whereas infusions during testing disrupted latent inhibition (**Figure 4A**). These experiments were simulated using a 20% reduction in inhibition (as per the learned irrelevance and blocking simulations) to mimic blockade of GABA-A receptors (see Methods). The model showed a similar pattern of results as observed by Piantadosi and Floresco (2014), with simulated GABA-A blockade increasing the latent inhibition effect if applied during conditioning, and eliminating latent inhibition if applied during testing (**Figure 4B**).

**Figure 4.**
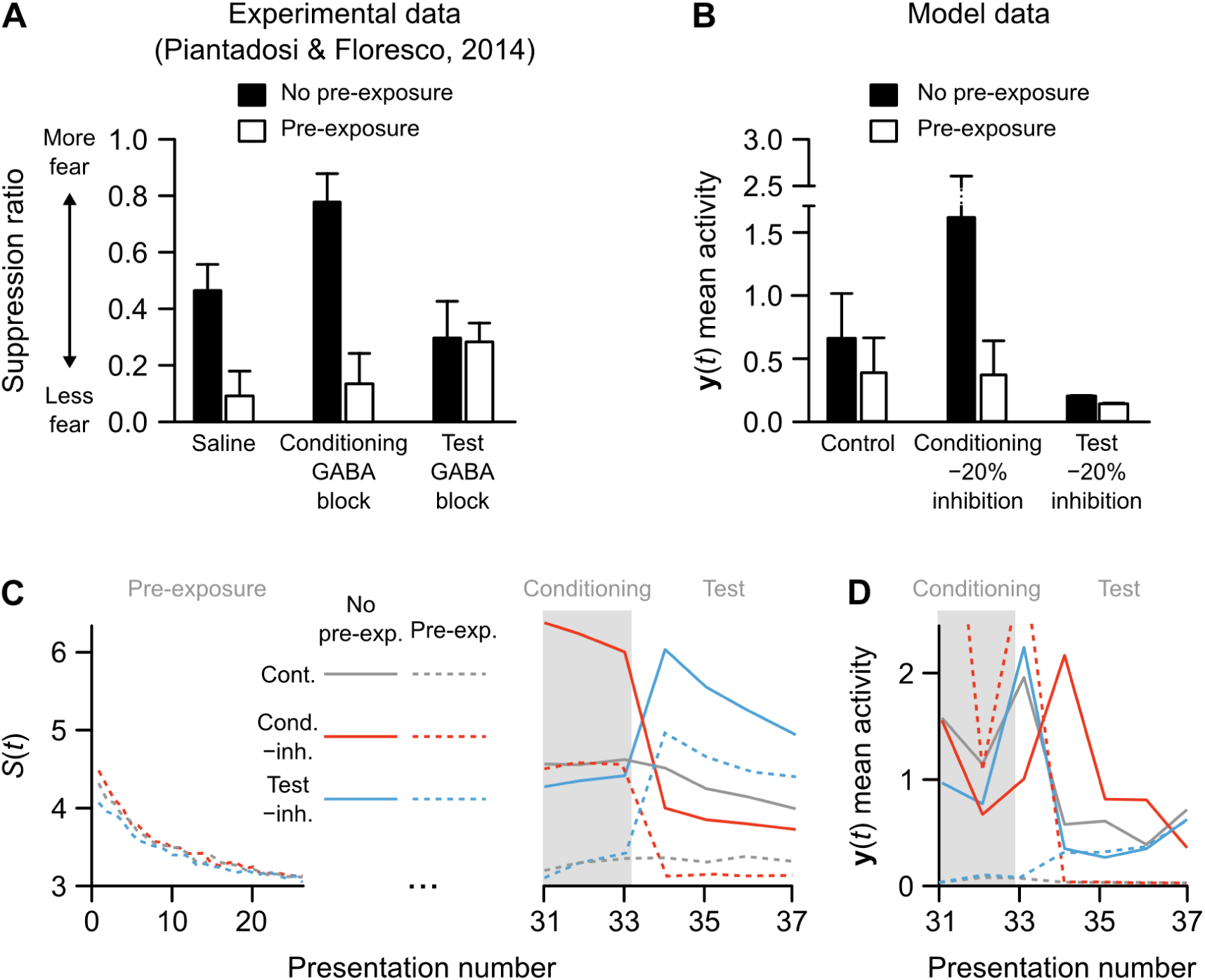
Simulation of experimental data on rodent latent inhibition. (A) Fear expression in rats in a latent inhibition paradigm in which animals were either pre-exposed (black bars) or not pre-exposed (white bars) to the conditioned stimulus, and treated with either saline, a GABA-antagonist during conditioning, or a GABA-antagonist during testing (reproduced from Piantadosi and Floresco (2014)). (B) Data from the model simulation of the same latent inhibition paradigm. Bars show median activity across 30 model runs (errorbars are 90% CI generated by bootstrapping 5-sample medians), of the average Amygdala layer activity during the final (test) stimulus presentations. (**C**) Cortex excitatory unit activity in response to stimuli across trials in an example run of the model. The downward curve during the first 30 presentations show learned irrelevance for all simulations in which the model was pre-exposed to the conditioned stimulus. (**D**) Amygdala activity levels in an example run of the model over trials with Conditioning and Testing epochs (as in the right-side panel of part C). Test period activity shows a pre-exposure effect in the control condition (solid versus dashed gray lines). This is amplified when inhibition is disrupted during conditioning (solid versus dashed red lines) but was lost when inhibition was disrupted during test (solid versus dashed blue lines).

The link between the salience signal (i.e., levels of excitatory unit activity) and Amygdala activity can be better understood by examining how each changed from one trial to the next (**Figures 4C-D**). As illustrated by the downward slope of activity among Cortex excitatory units over pre-exposure trials, the network receiving pre-exposures learned to ignore the CS over the preconditioning period, so that during the conditioning period (gray shaded area in **Figures 4C** and **4D**) Cortex activity was too low to always push the Amygdala past threshold, *θ*, making it less likely to trigger learning in the Amygdala competitive network (see equations (10)). The salience signal during conditioning for the non-pre-exposure tests of the model, however, resulted in a sufficiently strong Amygdala response to the CS to induce learning in the competitive network. When inhibitory signaling was experimentally reduced during conditioning, both Cortex activity and Amygdala learning were amplified. However, this learning included many excitatory units that were not coding the CS, so that the PE network remained relatively inactive during testing (**Figures 4C-D**). In contrast, when inhibitory signaling was reduced during testing, many Cortex units become active in both PE and NPE conditions. Since the Amygdala was modeled as a competitive network, the overactivity of non-CS coding inputs interfered with the activity of CS-coding inputs, resulting in overall low Amygdala responses in both PE and NPE conditions (**Figures 4C-D**). These results provide a new interpretation of the Piantadosi and Floresco (2014) data, one that that emphasizes the importance of the interaction between a relevance code in the mPFC and learning rates in a second system such as the amygdala.

### Inhibitory relevance-learning network with amygdala module recapitulates effect of optogenetic manipulations on fear behavior

The second simulation of experimental results addressed work by Courtin et al. (2014), which examined how the activity of PV+ interneurons in the mPFC controls fear expression. As described in Methods, PV+ interneurons are the cells modeled by the Cortex inhibitory unit, **I**(*t*). We simulated the experiments using the same network and parameters, including the Amygdala layer, as used to simulate latent inhibition above (see Methods). The original study by Courtin et al. (2014) demonstrated that optogenetic stimulation of PV+ interneurons in the mPFC resulted in increased freezing activity in mice, both before conditioning and, even more prominently, when stimulation was paired with a CS following extinction (**Figure 5A**, *left*). When we applied the same protocol on the model, using a reduction in inhibitory inputs to mimic optogenetic silencing (see Methods), the same pattern of activity was observed in the Amygdala layer (**Figure 5B**, *left*). The original experiments also showed that activation of mPFC PV+ interneurons decreased freezing to a conditioned CS (**Figure 5A**, *right*). This was consistent with activity patterns in the model in a subsequent set of simulations (**Figure 5B**, *right*). These results further support the hypothesis that the effects of the mPFC on efferent regions that control behavior is related to a potential relevance code mediated by feed-forward, divisive inhibition. They also offer evidence that the present model, in spite of its simplicity, captures an essential relationship between the role of inhibition in the mPFC region and the competitive network in the amygdala (Rashid et al., 2016).

**Figure 5.**
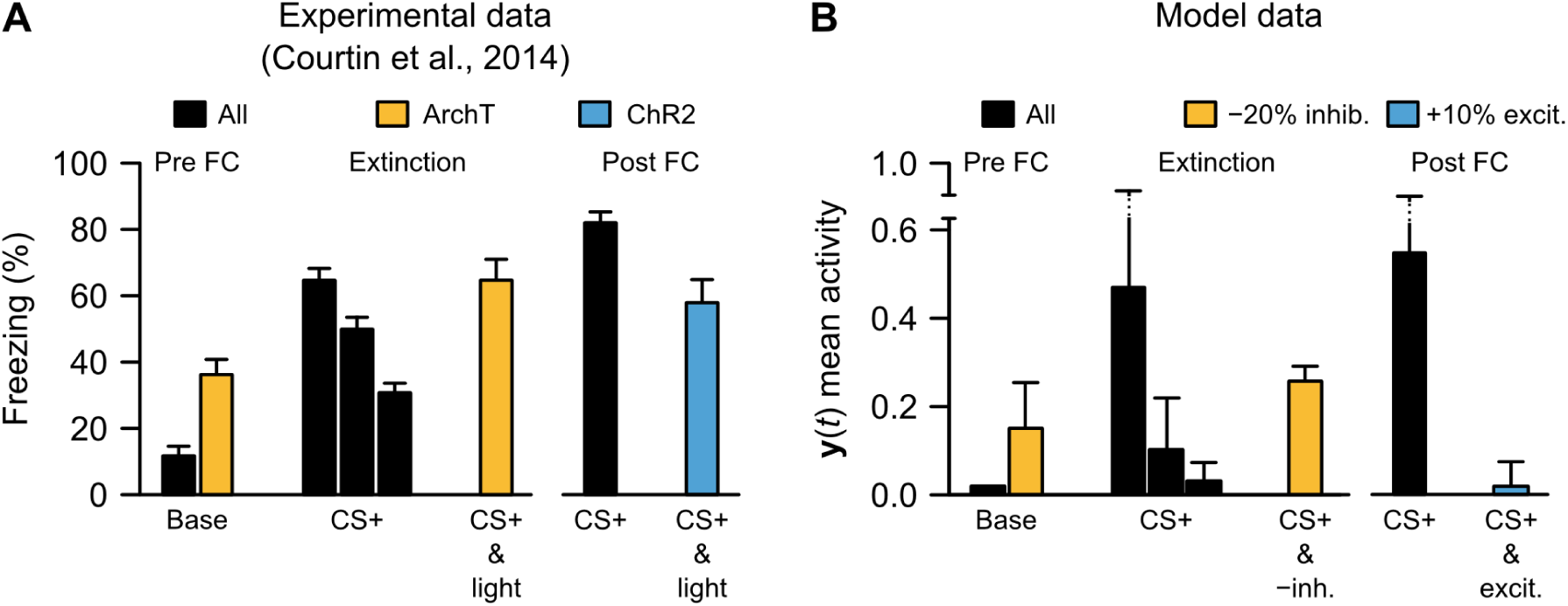
Simulation of experimental data on role of feed-forward inhibition in freezing behavior. (**A**) Experimental data illustrating the effects of optogenetic inhibition (“ArchT”) or excitation (“ChR2”) of the medial prefrontal cortex parvalbumin-expressing inhibitory neurons (modified from Courtin et al. (2014)). Inhibitory neuron inhibition was performed both before conditioning (”Base”) and following conditioning and extinction (CS+ & light). Inhibitory neuron excitation was performed following conditioning (right side, CS+ & light). (**B**) Replication of general patterns of inhbitory neuron manipulations (part (E)) in the model, substituting -20% inhibition for “ArchT” and +10% excitation (i.e., increased *W* ^**I**→**E**^ weights by 10%) for “ChR2”.

### Relevance learning can be multiplexed with input classification

A final set of simulations was used to investigate a key computational advantage to using inhibitory neuron plasticity for relevance learning, specifically by providing a method by which the brain could multiplex relevance learning with learning about other stimulus information. If the overall level of excitatory activity encodes relevance (via *S*(*t*)), and this is controlled by feedforward inhibition, then the excitatory synapses in the network should still be free to control the specific pattern of **E**(*t*) to encode other information. This can be described mathematically by viewing the excitatory Cortex activity patterns **E**(*t*) as vectors, where the norm (length) of the vector is a signal of relevance, but the position that the vector points in encodes other aspects of a stimulus, such as orientation, frequency, or category.

To test this idea, the network was trained to categorize 10 different stimulus classes, with only one of these paired with a reward. The prediction was that the network could learn information about relevance and also learn to respond with output patterns specific to the correct stimulus category. To train the network to categorize stimuli, we employed a softmax output layer (see Methods) and trained the excitatory pathway in the network with backpropagation of error (Rumelhart et al., 1986) (**Figure 6A**). It is worth noting that although backpropagation of error is traditionally not seen as a biologically realistic learning algorithm, there is evidence that it could be approximated with biologically realistic mechanisms (Scellier and Bengio, 2017; Lillicrap et al., 2016; Guergiuev et al., 2016; Zenke and Ganguli, 2017). Furthermore, independent of the specific algorithm used, the goal of the simulation was to offer a proof of the multiplexing concept. Training of the excitatory pathway with backpropagation was done concurrently with training of the feedforward inhibition pathway using the relevance learning algorithm (as described in Methods). Over the course of training, the network learned to dissociate the rewarded stimulus category from the unrewarded ones, via the relevance signal, *S*(*t*) (**Figure 6B**). Importantly, *S*(*t*) did not differentiate between the unrewarded categories (**Figure 6B**, *red lines*), demonstrating that it was not encoding the categories, *per se*, but their relevance for predicting reward. At the same time, the set of ’Category’ output units did learn to differentiate all 10 categories of stimuli. A cross-entropy loss function was used to evaluate the success of categorization, with lower values indicating a higher degree of separation between the categories. Over the course of 20 simulated seconds of training, this measure dropped to almost zero and the output layer was achieving roughly 95% accuracy on average (**Figure 6C**). Importantly, relevance learning and category learning were operating simultaneously. These results demonstrate the potential for multiplexing relevance signals with other stimulus information by using the overall level of excitatory activity as a code for value.

**Figure 6.**
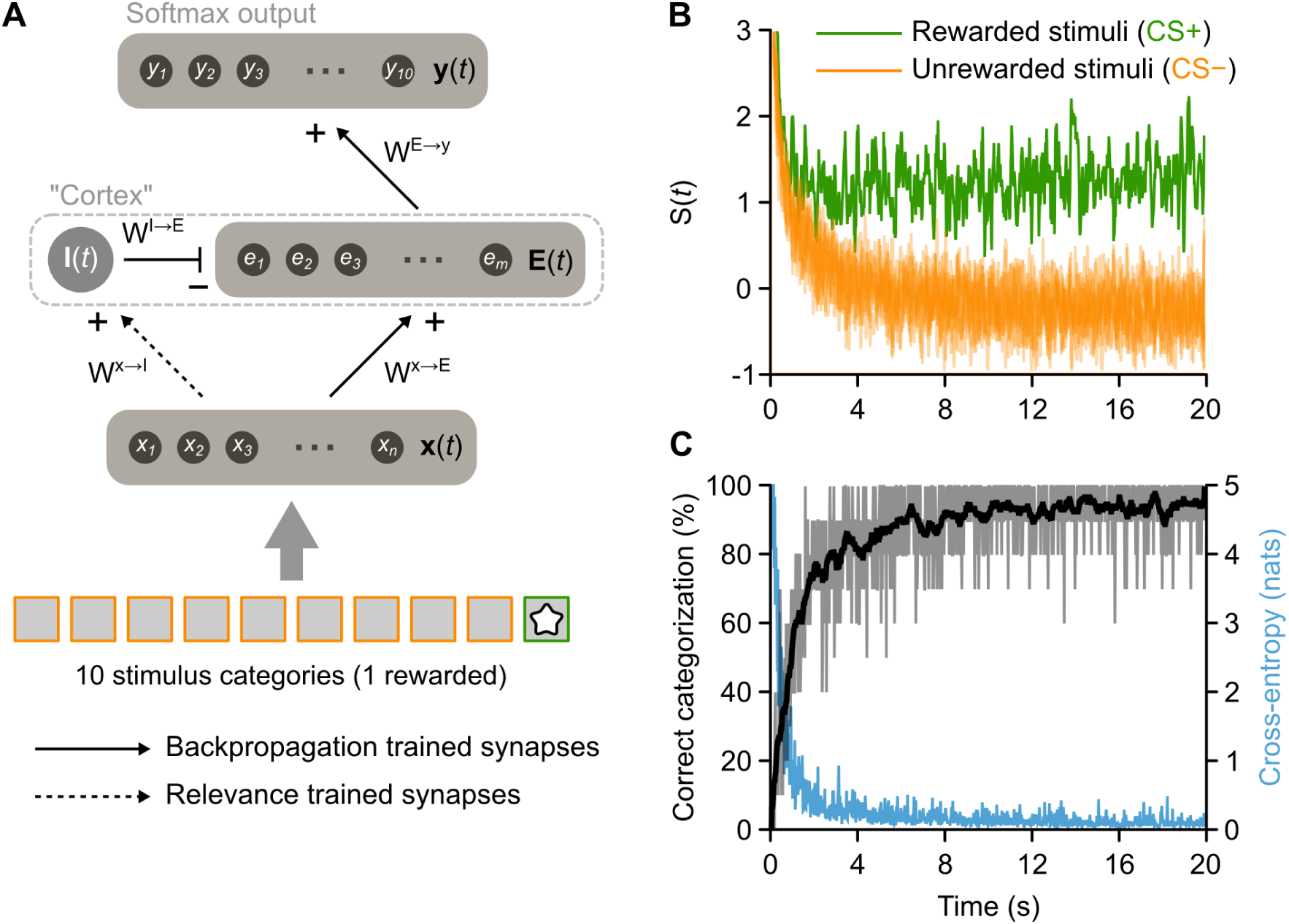
Multiplexed stimulus category and relevance codes via simultaneous excitatory and inhibitory learning. (**A**) Diagram illustrating modified model that included both the mechanisms described above for relevance learning (on *W* **^x→I^** synapses) in addition to mechanisms learning an output vector that matches categories presented as input (backpropagation algorithm applied to the *W* ^**E**→**y**^ and *W* ^**x**→**E**^ synapses). As illustrated by bottom boxes, only one of ten stimuli presented to the network was rewarded. (**B**) Average excitatory unit responses to the presented CS+ (red) and the nine CS-stimuli (blue) over trials. As in previous simulations, the network quickly learns respond more strongly to the CS+. (**C**) Performance of the model on input classification. Over the same trials that the network learns to discriminate CS+ and CS-stimuli, it also becomes capable of matching the output vector to the input. Black trace shows percent of presentations that stimuli are correctly classified, which increases quickly before reaching a plateau. Mauve trace shows the cross-entropy, an information measure (in natural units of information) based on the output activity distribution that is inversely related to the strength of input classification.

## Discussion

The simulations presented here explored the link between relevance assignment and neural disinhibition. They were motivated, in part, by the hypothesis that relevance learning impairments, much like those observed in schizophrenia, can be caused by a disruption to feed-forward inhibitory processes.The model was structured as simply as was necessary to examine this hypothesis, incorporating three core mechanisms. First, relevance was coded as the overall excitatory activity in the ‘Cortex’ layer. Specifically, the norm (length) of the excitatory units’ activity vector was treated as a reinforcement learning value function, though “unsigned” in that it treated positive (reward) and negative (punishment) values equivalently (Figure 1). Second, the model used feed-forward inhibition–i.e., the connections from the ‘Sensory’ input layer to Cortex inhibitory units–to control the overall level of Cortex excitatory activity. When paired with the first mechanism, the result was that disruptions to inhibition led to failures in relevance attribution (Figures 2 & 3). Third, the model used an established reinforcement learning algorithm, temporal difference learning, to train the feed-forward inhibitory connections and thereby learn to differentiate relevant versus irrelevant stimuli. When these mechanisms were further connected in sequence with a an output (the ‘Amygdala’) that used a competitive learning mechanism, this led to specific predictions about how disruptions to inhibition would alter fear behavior (Figure 4).

These three mechanisms are highly consistent with previous empirical work. The first mechanism, linking overall levels of excitation with a code for unsigned value, matches recent data on the importance of mPFC disinhibition for coding relevant situations (Pi et al., 2013; Courtin et al., 2014; Pinto and Dan, 2015; Kim et al., 2016a,b), including the observation that net levels of activity in putative pyramidal neurons increases near reward sites (Insel and Barnes, 2015). The second mechanism, assigning control of this relevance code to feed-forward inhibition, matches empirical findings on the importance of inhibition for behaviors like latent inhibition (e.g. (Piantadosi and Floresco, 2014)). It also matches decades of work linking relevance impairments in schizophrenia (Mcghie and Chapman, 1961; Venables, 1964; Lang and Buss, 1965; McGhie, 1970; Garmezy, 1977; Johnson, 1985; Kapur, 2003) with evidence that inhibitory neurons, and in particular classes of inhibitory neurons supporting feed-forward inhibition, may be differentially compromised in the disease (Benes and Berretta, 2001; Lewis et al., 2005; Lewis and Moghaddam, 2006; Lewis, k 2014; Gonzalez-Burgos et al., 2015; Krystal et al., 2017). Finally, the third mechanism, wherein inhibitory interneuron plasticity is the means for learning to differentiate relevant versus irrelevant stimuli, is consistent with findings that the neural connections supporting feed-forward inhibition are plastic (Kullmann et al., 2012; Kullmann and Lamsa, 2011; Bartos et al., 2011), in some cases requiring NMDA receptors with a well established importance for associative plasticity (Lamsa et al., 2007, 2005; Le Roux et al., 2013). There has been considerable recent interest in understanding the functions of inhibitory neuron plasticity (Barron et al.; Sprekeler; Hennequin et al.). These three mechanisms together comprise a more general theory of cortical coding: inhibitory plasticity may act in parallel with excitatory plasticity to accomplish different learning functions–while excitatory plasticity could provide a mechanism for learning various tasks, such as categorization, inhibitory plasticity could simultaneously provide a mechanism for relevance learning (Figure 5). This suggests a multiplexed code in the neocortex, and brings together the known specificity of inhibitory neuron plasticity with theories about their importance for maintaining EI balance.

As this was an abstract neural network model, many features of the real brain were absent. The most notable was the absence of feedback connections within Cortex. By excluding these connections, the mathematical complexity of calculating synaptic balances and their experience-dependent changes could be minimized, and it became possible to isolate those processes sufficiently to develop the learning algorithms that could explain the behavioral phenomena of interest. The results demonstrated that plasticity in the synapses connecting inputs to inhibitory neurons is sufficient to support relevance learning. Such a mechanism also causes relevance learning to be dysfunctional following disrupted inhibition. Other authors have taken a more biologically realistic approach to addressing the function of inhibitory neurons for learning and memory, including adding organized, feedback excitatory connections (Murray et al., 2014), along with organized, “motif” connections of specific inhibitory neuron types (Yang et al., 2016). The temporal precision of feed-forward inhibitory connections has also been proposed to regulate plasticity in excitatory neurons (). The findings from these more complex networks are consistent with those presently observed, and it seems likely that there is a meaningful mapping between the two. The present emphasis on simplicity for the sake of mathematical tractability offers only a first step toward understanding and treating disorders like schizophrenia.

In addition to excluding certain properties of the real brain, some other features of the model were kept abstract in order to remain agnostic as to their potential biological implementation. For example, the computed salience signal (*S*(*t*)) and corresponding prediction error responsible for maintaining EI balance (*β*(*t*)) might be integrated locally within the cortex itself (e.g. by feedback connections from excitatory to local inhibitory neurons), or computed by subcortical regions. As described in the Introduction, it is well known that networks maintain a strong balance between excitation and inhibition (Wehr and Zador, 2003; Zhang et al., 2003; Dorrn et al., 2010; Gabernet et al., 2005; Wilent and Contreras, 2005; Daw et al., 2007; Chittajallu and Isaac, 2010; Poo and Isaacson, 2009; Anderson et al., 2000; Xue et al., 2014; Haider et al., 2006) (see also the review by (Froemke, 2015)). While the process by which networks maintain this balance is not fully known, it has been proposed to be supported by plasticity at inhibitory synapses in response to feedback excitatory signals (Vogels et al., 2011; Luz and Shamir, 2012). Recent work has demonstrated local changes in synaptic scaling at both inhibitory and excitatory synapses following local changes in excitation (Barral and D Reyes, 2016). One possibility is that while the local network is capable of maintaining EI balance, the EI balance set-point can be adjusted by signals from extrinsic neuromodulators. Many neuromodulators are capable of signaling reinforcement or salience. While dopamine has received much of the attention in this regard, acetylcholine has also recently come under the spotlight (e.g. (Kuchibhotla et al., 2017; Letzkus et al., 2015, 2011; Hangya et al., 2015)). Disinhibition may also involve a particular class of inhibitory neurons that contain vasoactive intestinal polypeptide (VIP+ interneurons (Pi et al., 2013)). Although the present model does not identify the specific biological process responsible for transmitting the salience and prediction error signals to inhibitory neuron synapses, it does predict that the plasticity processes responsible for relevance learning directly impact the mechanisms for maintaining EI balance; otherwise, EI balance maintenance would be constantly working to compensate for the adjustments to EI balance associated with relevance learning. To accurately capture the mechanisms of the supervisory process, it will likely be necessary to increase the complexity of the model by including feedback connections between excitatory neurons, and connections from excitatory to inhibitory neurons.

The primary prediction of the present work is that relevance learning involves plasticity on the feed-forward, input-to-inhibitory neuron synapses in a cortical circuit. This prediction is consistent with recent data showing that the number of excitatory synapses onto parvalbumin-expressing inhibitory neurons is reduced in schizophrenia (Chung et al., 2016). It also contributes to an idea gaining momentum that during memory storage patterns of inhibitory modifications mirror excitatory modifications, in order to maintain EI balance and reduce inappropriate recall, i.e. that there is an “inhibitory engram” (Barron et al.). The present model has many parallels with this idea. Specifically, inhibitory synapses in the model learned to match any increases in excitatory synapses in order to keep the ‘Cortex’ output at the homeostatic set-point. Only when the prediction error signal indicated that an input was relevant was this mirrored inhibition relaxed to allow the excitatory activity to increase.

The simulations also addressed how the proposed code for relevance might impact learning and activity in a post-synaptic region. With the aim of simulating experimental fear-learning data, the post-synaptic region chosen was the amygdala, which was modeled as a competitive learning network (Rumelhart and Zipser, 1985). Use of a competitive network is consistent with known properties of the mammalian amygdala (Yiu et al., 2014; Rashid et al., 2016). By combining our proposed relevance code at one layer of the model with a competitive learning rule at the next, the model was capable of responding selectively to only those specific stimuli that had been paired with reinforcement signals in the past. The model thereby became capable of replicating behavioral patterns from both pharmacological (Piantadosi and Floresco, 2014) and optogenetic (Courtin et al., 2014) manipulations to mPFC inhibitory neurons.

One area that was not explored in the current set of experiments was the increasingly apparent link between pathologies of EI balance and deficits in social behavior and motivation (Rubenstein and Merzenich, 2003; Yizhar et al., 2011; Nelson and Valakh, 2015). It seems likely, however, that some of the more basic results from the present investigations could offer more understanding for why some individuals have more difficulty filtering or dynamically processing social information. Tackling these complex problems will require a convergence of multiple experimental and theoretical approaches, and mathematically tractable network models that include excitatory-inhibitory interactions will be an essential tool.

Altogether, our theoretical investigations provide a potential explanation for why behaviors such as gating and relevance learning may emerge from disrupted inhibition, and therefore, how pathologies of inhibition may underlie neuropsychiatric conditions such as schizophrenia. The ideas are not entirely new, in many ways they reformulate a long existing hypothesis that schizophrenia is a disruption of feed-forward inhibition (Johnson, 1985). But the model offers a computational description of the process with defined links between several functional elements. Furthermore, it offers valuable predictions about the importance of plasticity in both excitatory and inhibitory neurons, lending insights into the normally functioning brain.

## Acknowledgements

We would like to thank Adam Santoro for his comments on an earlier draft of this manuscript. This work was supported by the Natural Sciences and Engineering Research Council of Canada (Discovery Grant RGPIN-04947), Google (Faculty Research Award 2016), and the Canadian Institute for Advances Research (Learning in Machines and Brains Fellowship).

